# Serotonin deficiency from constitutive SKN-1 activation drives pathogen apathy

**DOI:** 10.1101/2024.02.10.579755

**Authors:** Tripti Nair, Brandy A. Weathers, Nicole L. Stuhr, James D. Nhan, Sean P. Curran

## Abstract

When an organism encounters a pathogen, the host innate immune system activates to defend against pathogen colonization and toxic xenobiotics produced. *C. elegans* employ multiple defense systems to ensure survival when exposed to *Pseudomonas aeruginosa* including activation of the cytoprotective transcription factor SKN-1/NRF2. Although wildtype *C. elegans* quickly learn to avoid pathogens, here we describe a peculiar apathy-like behavior towards PA14 in animals with constitutive activation of SKN-1, whereby animals choose not to leave and continue to feed on the pathogen even when a non-pathogenic and healthspan-promoting food option is available. Although lacking the urgency to escape the infectious environment, animals with constitutive SKN-1 activity are not oblivious to the presence of the pathogen and display the typical pathogen-induced intestinal distension and eventual demise. SKN-1 activation, specifically in neurons and intestinal tissues, orchestrates a unique transcriptional program which leads to defects in serotonin signaling that is required from both neurons and non-neuronal tissues. Serotonin depletion from SKN-1 activation limits pathogen defense capacity, drives the pathogen-associated apathy behaviors and induces a synthetic sensitivity to selective serotonin reuptake inhibitors. Taken together, our work reveals new insights into how animals perceive environmental pathogens and subsequently alter behavior and cellular programs to promote survival.

**KEY POINTS:**

- Identify an apathy-like behavioral response for pathogens resulting from the constitutive activation of the cytoprotective transcription factor SKN-1.
- Uncover the obligate role for serotonin synthesis in both neuronal and non-neuronal cells for the apathy-like state and ability of serotonin treatment to restore normal behaviors.
- Characterize the timing and tissue specificity of SKN-1 nuclear localization in neurons and intestinal cells in response to pathogen exposure.
- Define the unique and context-specific transcriptional signatures of animals with constitutive SKN-1 activation when exposed to pathogenic environments.
- Reveal necessity for both neuronal and non-neuronal serotonin signaling in host survival from pathogen infection.

## INTRODUCTION

Avoidance of a potential pathogen is one of the earliest defenses an organism employs to ensure survival. In its natural environment, *Caenorhabditis elegans* feed upon a variety of bacterial diets, with varying dietary benefits and pathogenic quality, which can affect organismal health [1, 2]. When exposed to a pathogenic microbe, *C. elegans* must initiate a proper immune response to defend against infection [3]. Even though *C. elegans* lack a classical adaptive immune system, they initiate a potent innate immune response, through the upregulation of genes like c-type lectins, anti-microbial peptides, and other defense mechanisms in order to fight off pathogenic infection [4].

*Pseudomonas aeruginosa* (PA14) is a human opportunistic pathogen that is lethal to *C. elegans* with prolonged exposure. The p38 mitogen activated protein kinase (MAPK) pathway and many of the downstream transcriptional effectors are evolutionarily conserved, indicating a critical role potentiating cellular responses. In *C. elegans*, the role of the p38 MAPK pathway in modulating the innate immunity system was revealed through genetic screens for sensitivity to pathogen infection; these include: *nsy-1*, a MAP kinase kinase kinase (MAPKKK) [5], *sek-1*, a mitogen-activated protein kinase kinase (MAPKK) [6], and *pmk-*1 MAP kinase (MAPK) [5]. The p38 pathway modulates the expression of downstream immune effector molecules (e.g., lysozymes, antimicrobial peptides, and C-type lectins) [7] to help ensure survival upon encountering pathogenic environments.

Among the physiological consequences of PA14 exposure is the rapid depletion of stored intracellular lipids in the intestine; a phenotype that manifests even before pathogen colonization and intestinal bloating can be measured [8].This striking phenotype of pathogen exposure is regulated by the multifaceted cytoprotective transcription factor SKN-1/NRF2, which canonically regulates phase 2 detoxification and redox balance [9, 10]. More recently, it has been studied for its essential role in the immune response [8, 11]. Upon activation, SKN-1 activates the expression of the phase II cellular detoxification system and multiple pathogen response genes [8, 11–13] that help protect the organism from pathogen exposure. Relatedly, animals with constitutive activation of SKN-1 display enhanced resistance to killing by pathogen exposure [8]. However, the enhanced innate immune response to pathogens occurs at the expense of organismal lipid homeostasis and results in a redistribution of lipid stores from the soma to germline, leading to impaired organismal health, oxidative stress resistance, and a shortened lifespan [8, 14]. Transcriptional redirection of activated SKN-1 away from pathogen resistance and immune response genes towards oxidative stress and xenobiotic detoxification genes restores metabolic homeostasis and prevents the depletion of somatic fat from the intestines thereby improving healthspan and lifespan [8]. As such, although SKN-1 activation plays a critical role in balancing cytoprotection, failure to turn off this response is pleiotropic.

Another mechanism *C. elegans* utilize to prevent lasting harm and death from pathogens is an avoidance phenotype [15–17]. Initially, naïve worms that have never experienced a pathogenic infection will preferentially move towards PA14 over *E. coli* through stimulation of olfactory behavioral responses [15, 18]. However, this preference does not persist as sophisticated intracellular and cell non-autonomous signaling pathways are initiated to switch the behavior of the host toward avoiding the pathogenic bacteria [19–21]. This brief exposure to PA14 functions as an important learning experience for the animal to avoid future exposure to the pathogen [22, 23] and this learned behavior to avoid pathogens can be transmitted to progeny as well [24].

[1]. The ability to avoid pathogenic bacteria, even after a brief exposure, is essential for the health of the organism, as the inability to leave can lead to a more rapid death. The signaling pathways driving aversion behaviors are complex, but clearly require neuropeptide signaling and downstream neurogenic targets [20, 23, 25]. Although intestinal bloating has been shown to drive leaving behaviors, several of the molecular details that influence the decision to leave pathogenic bacteria for other food sources is not fully understood. Among the array of neurometabolic signaling molecules that mediate homeostasis, serotonin plays a critical role in the modulation of gut-neuronal signaling in response to dietary inputs and regulating immune responses [26]. Serotonin synthesis, in response to environmental and nutritional cues, regulates fat mobilization, which may be utilized to elicit immune responses [27]. Here we demonstrate that animals with constitutive activation of SKN-1, without the capacity to turn it off, have an activated innate immune system yet fail to initiate leaving behaviors from PA14, despite ultimately dying from exposure. This apathetic early response to pathogen is driven by cell non-autonomous communication between the digestive and nervous systems stemming from defective serotonin signaling. Understanding why constitutive SKN-1 activation leads to deficiencies in pathogen leaving behavior will enhance our understanding of the molecular drivers of pathogen avoidance and the role that SKN-1 transcriptional activation plays in this essential survival response to environmental pathogens.

## RESULTS

### Constitutive activation of SKN-1 drives pathogen apathy

Studies of host responses to pathogenic diets traditionally use preference plates where animals are placed equidistant from two different food options (**Figure 1A** and Table S1) and initial choice between the two options are recorded. Beyond this initial choice, animals can choose to remain on one food source or explore other options, but the decision matrix used in this complex behavior required additional study. Despite the numerous reports on pathogen food choice, there has not been a consistent experimental design for these studies and as such, we tested several paradigms of PA14 culturing to standardize our studies of leaving index. Ultimately, our studies showed the most dynamic leaving response when the PA14 pathogen was cultured at 37°C for 24 hours, followed by 48 hours at 25°C [28] (Figure S1A). We compared the choice dynamics of WT and *skn-1gf* mutants with constitutive SKN-1 activity and discovered that although initial choice was not significantly different, *skn-1gf* mutants that first choose the PA14 bacteria failed to leave the pathogen environment like age-matched WT animals (**Figure 1B** and Figures S1B-D); behaving similar to animals given a choice of *E. coli* or a non-virulent PA14 strain harboring a mutation in the *gacA* gene (Figure S1E-F) [29]. This difference in choice was not influenced by animal movement speed, as all animals reach the first bacteria at similar rates (Figure S1G), nor did minor differences in developmental timing in the *skn-1gf* mutant influence choice kinetics (Figure S1H).

**Figure 1.**
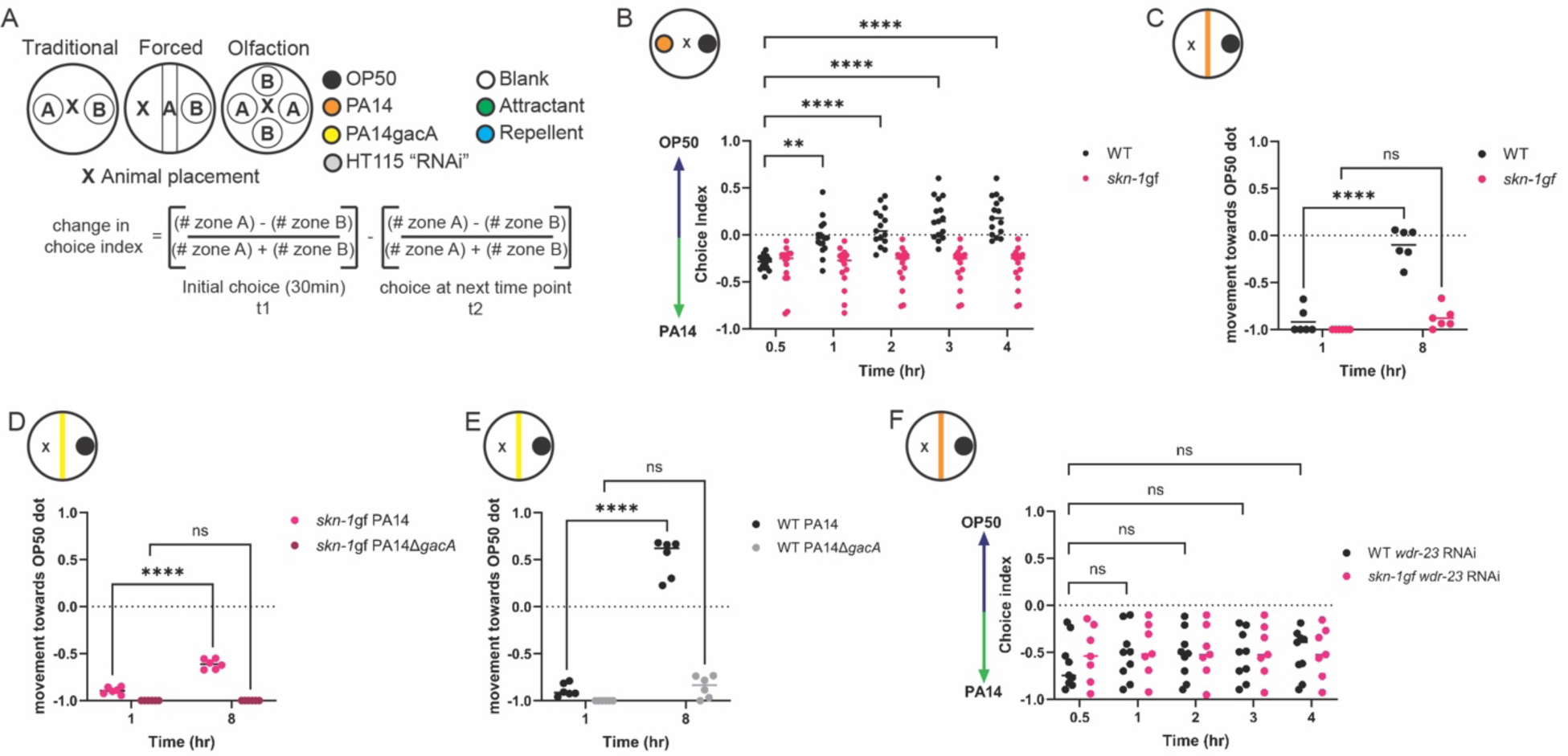
Activation of SKN-1 drives pathogen apathy. (**A**) Graphical representation of the variations in food choice assays performed. (**B-C**) Apathy displayed by *skn-1gf* worms in the traditional choice assay (**B**) or forced choice assays (**C**). The apathy response of *skn-1gf* worms is attenuated in the non-pathogenic strain PA14*ΔgacA* (**D**) and resembles the leaving behavior of WT animals (**E**). Both WT and *skn-*1*gf* mutant animals display apathy for PA14 following depletion of *wdr-23* by RNAi (**F**). Each of the food choice assays comprised of N≥3 and analyzed via two-way ANOVA test; **(p<0.01) ***(p<0.001) ****(p<0.0001).

A confounding factor of traditional food choice assays is the variation in the initial choice at the start of the experiment, which cannot be overcome by changing the distance between the two diets (Figure S1I) and limits the power to study leaving behavior after the initial choice. To overcome this limitation, we developed a “forced” choice assay (**Figure 1A**) where animals must interact with one diet first, as a line on the plate, and can then choose to leave that food source for a second food option. In this experimental set up, *skn-1gf* mutant animals reach the PA14 line within 60 min (Figure S1J), similar to WT animals, but remain on the pathogen, whereas the majority of WT animals move toward the *E. coli* option over an 8-hour time course (**Figure 1C**); and this phenotype is sensitive to the pathogenicity of PA14 as neither genotype choses to leave the attenuated PA14*ΔgacA* mutant (**Figure 1D-E**). We tested whether other mechanisms of SKN-1 activation could impede pathogen leaving and confirmed that RNAi knockdown of the *wdr-23* gene, a negative regulator of SKN-1 activity that results in potent activation of SKN-1 when impaired [30, 31], was sufficient to delay WT animals from leaving the pathogen food source (**Figure 1F**) and was not additive with *skn-1gf* mutation (Figure S1K-L). This result supports a model where constitutive SKN-1 activation impedes leaving behavior, but also uncouples the behavioral response from the SKN-1-dependent and pathogen-related phenotype of somatic lipid depletion that is suppressed by the *E. coli*/K-12 “HT115” diet used for RNAi [8]. Taken together, we coin this failure to leave a pathogenic environment as a form of “pathogen apathy”.

### Pathogen apathy is a specific neuronal response

SKN-1 has established roles in ASI neurons and most recently we discovered that the phenotypes associated with the *skn-1gf* allele begin in the nervous system to initiate systemic responses cell non-autonomously [32]. With this in mind, and in light of the sensory role of the ASI neurons, we first tested whether *skn-1gf* mutants are generally defective in olfaction, which could influence their decision to not move off the pathogenic food and found that neither chemoattraction to diacetyl or isoamyl alcohol (**Figure 2A**) or chemorepulsion from octanol or nonanone (Figure S2A) were impaired. The avoidance of a pathogenic environment is a learned behavior and informed by past experience. To determine if animals with constitutive activation of SKN-1 could learn to avoid pathogen, as previously reported in WT animals that are subjected to short exposures to the pathogen as a training paradigm [33, 34], we trained age-matched WT and *skn-1gf* mutants through pre-exposure to PA14 for 2 and 4 hours before being subjected to a traditional food choice assay plate (**Figure 2B**), which as expected resulted in enhanced avoidance of PA14 the 30min to 2 hours experimental time course for WT animals (**Figure 2C** and Figure S2B-C). In contrast, neither 2 hours or 4 hours of training was sufficient to change the apathy behavior of the *skn-1gf* mutants (**Figure 2D** and Figure S2D-E). Taken together these findings suggests that the behavior defect that induces apathetic leaving behaviors is a specific choice, but not clearly associated with the perception of volatile odorants within the diet.

**Figure 2.**
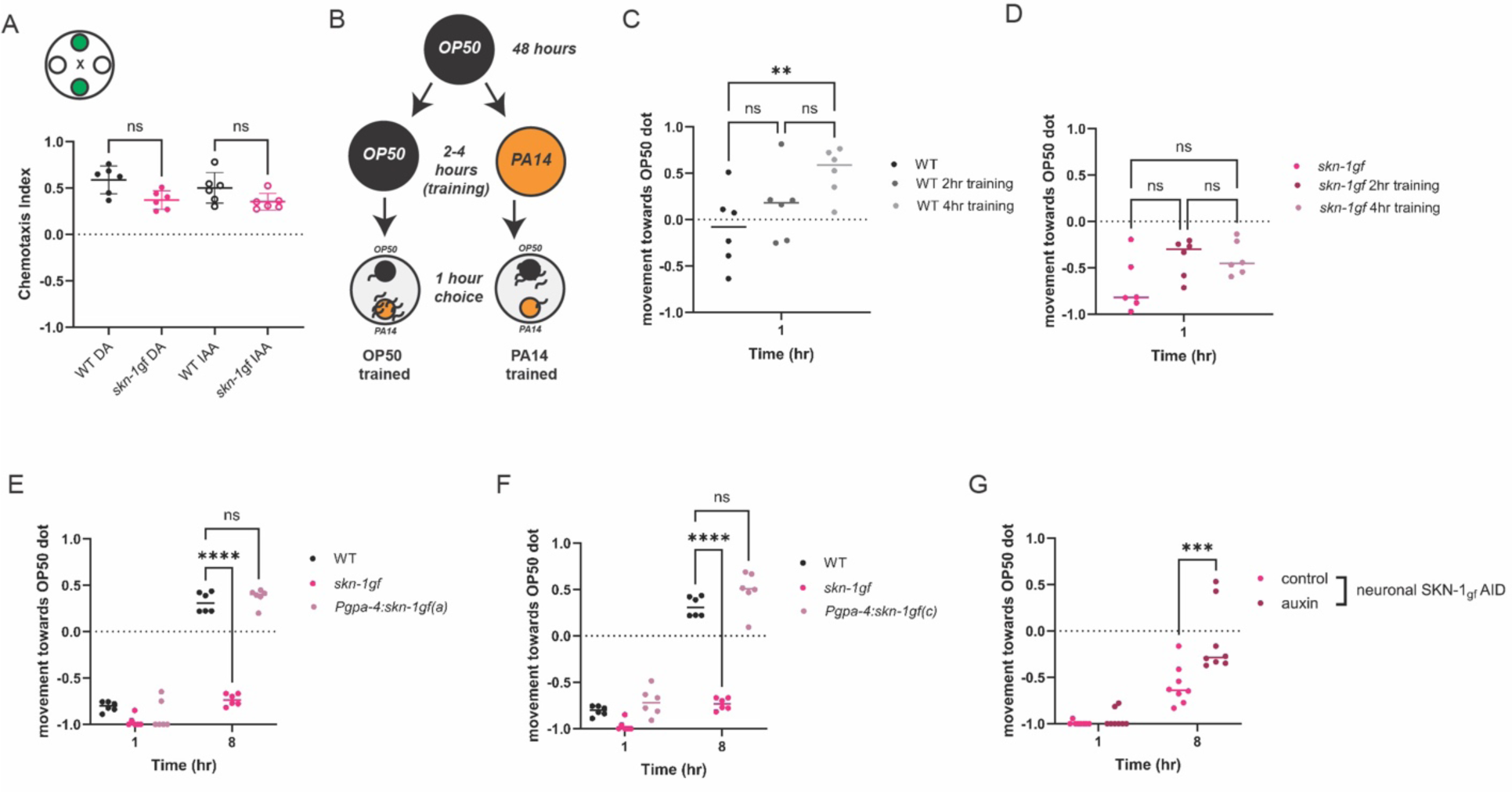
Pathogen apathy requires neuronal SKN-1. Chemotaxis towards (**A**) attractants [diacetyl (DA) and isoamyl alcohol (IAA)] are similar for both WT and *skn-1gf* worms. (**B**) Graphical representation of pathogen training and subsequent food choice assay. Differences in pathogen leaving response in (**C**) WT and (**D**) *skn-1gf* worms following pathogen training for 0hr, 2hr, and 4hr of exposure. (**E**) Pan-neuronal degradation of SKN-1gf restores pathogen leaving behavior in the *skn-1gf* mutant. Specific expression of SKN-1gf isoform a (**F**) or isoform c (**G**) in ASI neurons does not elicit failed pathogen leaving behaviors as observed in the *skn-1gf* mutant. Each of the chemotaxis assay comprised of N≥3 and analyzed via one-way ANOVA test. Each of the food choice assay comprised of N≥3 and analyzed via two-way ANOVA test; **(p<0.01) ***(p<0.001) ****(p<0.0001).

Our previous work has demonstrated that SKN-1 activation specifically in ASI neurons is sufficient to drive cell non-autonomous effects throughout the organism [32]. With this in mind, we next expressed the two *skn-1* isoforms impacted by the *skn-1gf* mutation exclusively in ASI neurons. Surprisingly, the expression of *skn-1a* (**Figure 2E**) or *skn-1c* (**Figure 2F**) could not replicate the failed pathogen leaving behavior observed in the *skn-1gf* mutant. We next examined if SKN-1 activity in neurons, in general, was necessary for the apathy behavior observed in the *skn-1gf* mutant by degrading the SKN-1gf protein, specifically in the nervous system, by an auxin-inducible degradation (AID) system [35]. Pan-neuronal degradation of SKN-1gf was sufficient to restore pathogen leaving behavior in the *skn-1gf* mutant (**Figure 2G**) and had no effect on the pathogen leaving behavior of WT animals with loss of SKN-1wt-AID in the nervous system (Figure S2F). Based on the necessity of SKN-1gf activity in the nervous system, but perhaps outside of the ASI neuron pair, we modified the expression of the *skn-1gf* isoforms pan-neuronally, but similar to ASI-specific expression, pan-neuronal expression could not phenocopy the *skn-1gf* mutant failure to leave the PA14 food source (Figure S2G-H). Collectively, these data reveal that *skn-1gf* activity in neurons is necessary, but not sufficient, to drive apathetic leaving behavior in response to pathogens and indicate that additional factors beyond neuronal expression are important.

### SKN-1 activation does not impact luminal distention in response to Pseudomonas

In response to stress, SKN-1 is stabilized and accumulates within the nucleus, particularly in the intestine [10, 32], where it can promote the transcription of cytoprotective genes that function to alleviate the stressful conditions [36–38]. Although *skn-1gf* mutants have constitutive activation of the SKN-1 protein, it is still rapidly turned over in the absence of exogenous stress [32]. To assess how pathogen exposure might differentially impact SKN-1 stabilization, we exposed animals harboring a CRISPR/Cas9-edited GFP at the C-terminal end of SKN-1wt or SKN-1gf to PA14 to assess protein stabilization and nuclear localization, which is easily observed in the relatively large cells of the *C. elegans* intestine. Outside the constitutive expression of both SKN-1wt and SKN-1gf in the ASI neurons [32], when exposed to PA14, SKN-1gf-GFP accumulated in the first two intestinal nuclei earlier than SKN-1wt-GFP (**Figure 3A-E**, Figure S3A-F and Table S2) and the abundance of nuclear-localized protein was greater in SKN-1gf-GFP mutants (**Figure 3F**). Because SKN-1gf accumulation in the nucleus is accelerated in response to PA14, these data suggest that the failure to leave a pathogenic environment is not a result of failed stabilization and subcellular localization of SKN-1.

**Figure 3.**
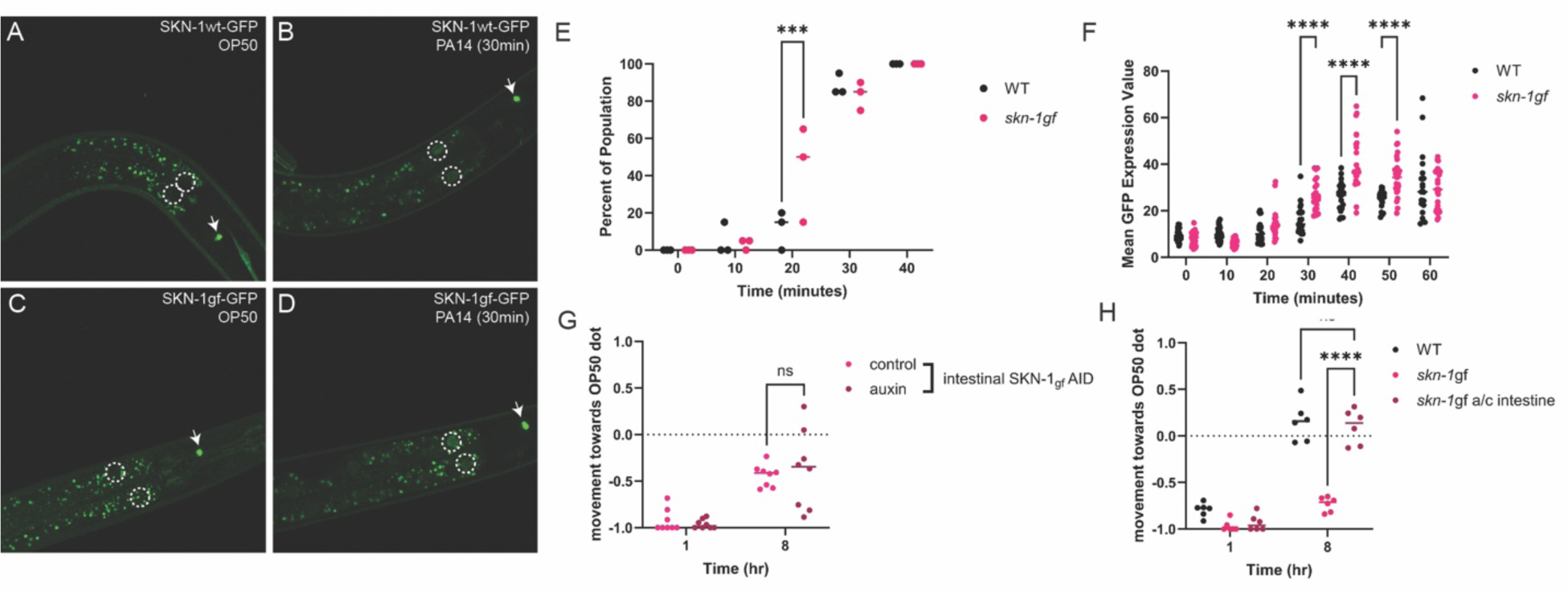
Precocious activation of intestinal SKN-1 in responses to PA14. (**A-D**) Intestinal accumulation and stabilization of SKN-1-gf-GFP and SKN-1wt-GFP upon PA14 exposure in comparison to OP50 (time point: 30 mins); the first two intestinal nuclei of each worm are outlined by white-dotted circles. Quantification of the percent of population (**E**) and intensity of nuclear localization (**F**) over the time course of PA14 exposure. (**G**) Intestine-specific degradation of SKN-1gf does not restore pathogen leaving behavior in *skn-1gf* mutant. (**H**) Co-expression of a and c isoforms of SKN-1gf in the intestine does not elicit failed pathogen leaving behavior, as observed in the *skn-1gf* mutant. Each of the nuclear accumulation time points comprised of N≥3 and analyzed via two-way ANOVA test; **(p<0.01) ***(p<0.001) ****(p<0.0001).

We next tested if intestinal expression of the *skn-1gf* allele was important for the failed pathogen leaving behavior of *skn-1gf* mutant by specifically degrading SKN-1gf-AID in the intestine. Activation of AID in the intestine had minimal capacity to restore pathogen leaving behavior in the *skn-1gf* mutant (**Figure 3G**) and no effect on SKN-1wt-AID animals (Figure S3G). Moreover, intestinal-specific co-expression of the gain-of-function variant of *skn-1a/c* (**Figure 3H**), nor the expression of the individual gain-of-function variants of *skn-1a* (Figure S3H) *or skn-1c* (Figure S3I), could lead to an impairment of pathogen leaving behavior. Taken together, these data reveal that although the SKN-1gf protein can be stabilized more rapidly within the intestinal nuclei of PA14-exposed animals, unlike the nervous system where SKN-1gf is required for pathogen apathy behaviors, expression within the intestine is neither necessary nor sufficient to influence pathogen leaving behavior.

Intestinal bloating in response to pathogen colonization in the gut is an important driver of pathogen leaving behaviors [19]. With this in mind, we measured intestinal lumen distention following exposure to PA14 and discovered that bloating was unremarkable between WT and *skn-1gf* mutant animals (Figure S3 J-M). As such, the failure to leave the PA14 diet is not due to the absence of the intestinal bloating trigger and also noting that WT and *skn-1gf* mutant animals display normal luminal morphology in the presence of *E. coli* OP50 bacteria. RNAi of *aex-5* or *egl-8* have previously been demonstrated induce intestinal bloating and pathogen leaving responses in WT animals [19]. However, only *egl-8* RNAi was capable of marginally inducing leaving behavior of the *skn-1gf* mutant animals, albeit not to the level of WT animals with normal *egl-*8 expression and far worse than WT animals subjected to *egl-8* RNAi (Figures S3 N-O). These data confirm that the distention of the intestinal lumen is unremarkable in *skn-1gf* mutant animals and the failure to leave the pathogen environment following the ingestion of PA14 is a defect in a downstream response.

### Unique and context specific transcriptional signature of SKN-1 activation and pathogen exposure

The sustained intestinal distension in response to PA14 exposure in animals with the *skn-1gf* allele when combined with the precocious stabilization and nuclear localization of SKN-1gf protein suggested to us that the pathogen apathy observed is a failure in responses following the initial exposure to pathogen. As such, to elucidate the molecular basis underlying the difference in pathogen leaving between WT and *skn-1gf* animals, we employed RNAseq to define differential transcriptional responses to PA14 exposure in animals with and without constitutively activated SKN-1 (**Figure 4A** and Table S3). We used the DESeq2 package in R to compare changes in transcriptional response across genotypes (WT or *skn-1gf*) and conditions (OP50 or PA14 exposure). Principal component analysis of the four sample groups (WT - OP50, WT - PA14, *skn-1gf* - OP50, and *skn-1gf* - PA14) clearly demonstrates that each sample is unique due to the lack of overlaps between groups (**Figure 4B**). Pairwise analysis of the four sample groups identified a significant number of genes both down- and up-regulated, along with a number of specific pathways altered, most of which were associated with metabolism and stress responses (**Figure 4A**). Although distinctly different, the analysis revealed similarities when conducting pairwise comparisons, three-way comparisons, along with identifying 1348 genes that are differentially expressed across all comparisons (**Figure 4C**).

**Figure 4.**
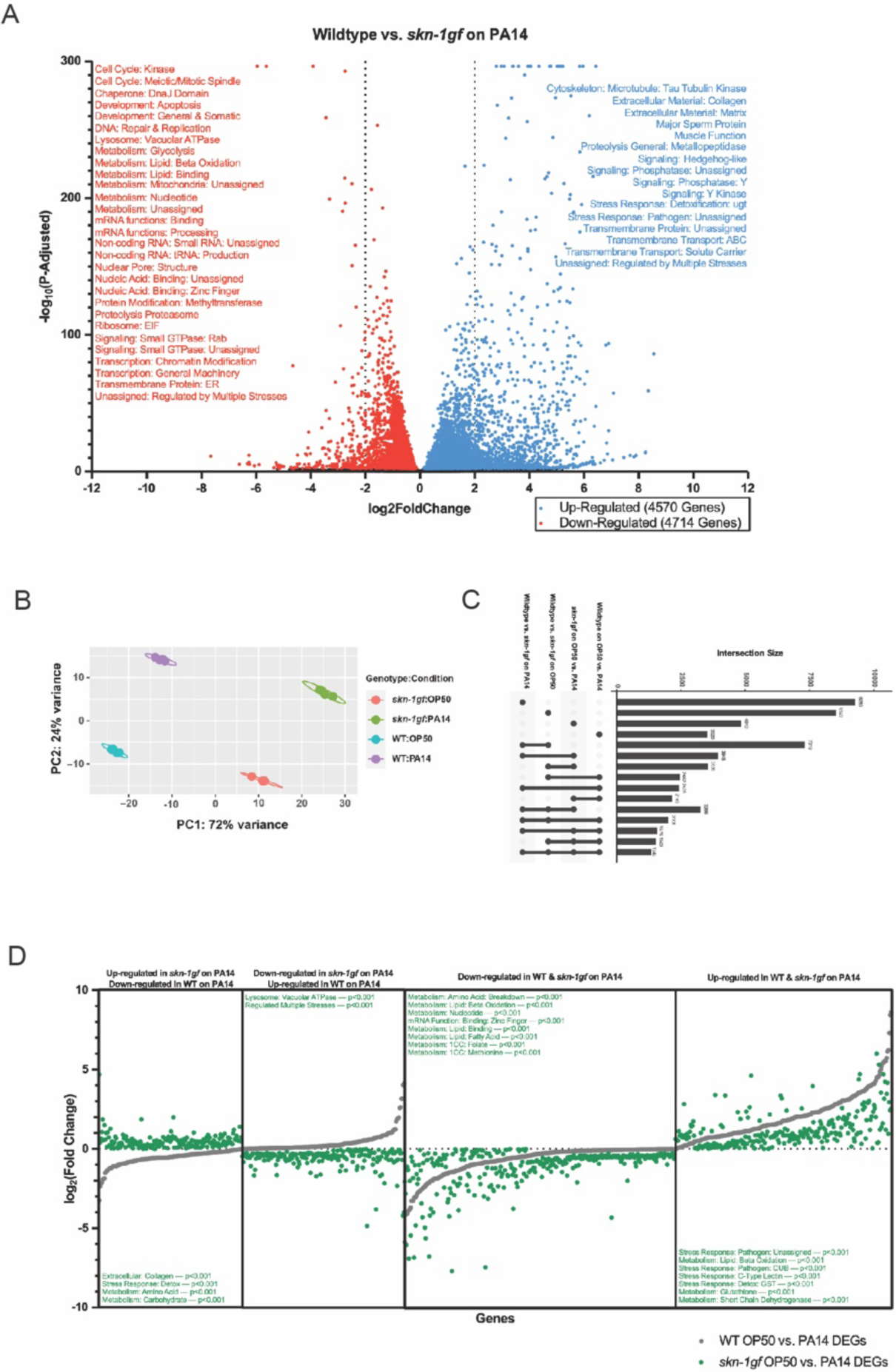
Context dependent transcriptional signature of PA14 exposure. (**A**) RNAseq of differential transcriptional responses to PA14 exposure in WT and *skn-1gf* animal (WT vs *skn-1gf* up-regulated in blue to the right and down-regulated in red to the left). (**B**) Principal component analysis of the four sample groups (WT - OP50, WT - PA14, *skn-1gf* - OP50, and *skn-1gf* - PA14) demonstrating lack of overlaps between groups, indicative of the uniqueness in the transcriptional responses. (**C**) Analyses of the various classes of genes with unique transcriptional response revealed similarities upon pairwise comparisons, three-way comparisons, along with identifying 1348 genes that are differentially expressed across comparisons. (**D**) Genotype-specific responses to PA14 identified 363 genes with reversed directionality of transcriptional response between WT and *skn-1*gf animals (170 down-regulated and 193 up-regulated in WT in comparison to *skn-1gf*). 579 genes with similar directionality but differential magnitude of the transcriptional response between WT and *skn-1gf* mutants (322 down-regulated and 257 up-regulated).

When examining genotype-specific responses to PA14 at the L4 stage, we identified 363 genes where the directionality of the transcriptional response was reversed; 193 that are up in WT but down in *skn-1gf* (**Figure 4D** and Figure S4A and Table S3) and 170 down in WT but up in *skn-1gf* (**Figure 4D** and Figure S4B and Table S3). Moreover, we identified 579 genes where the directionality was the same, but the magnitude of the transcriptional response was significantly different between WT and *skn-1gf* mutants; 322 down-regulated (**Figure 4D** and Table S3) and 257 up-regulated (**Figure 4D** and Table S3). Intriguingly, gene set enrichment analysis [39] revealed reproductive systems, the pharynx, and socket cells, which make the pore where sensory dendrites extend into the cuticle and exterior were over represented in the context-dependent transcriptional response (Figure S4C). The phenotypes associated with the differentially regulated genes included intestinal function, shortened lifespan, and behavior (Figure S4D), and gene ontology terms that were enriched included adherens junctions, proton transport across membranes, and biosynthetic and catabolic processes (Figure S4E). Taken together, these data reveal the unique and context-specific transcriptional signatures of animals with constitutive SKN-1 activation when exposed to different bacterial environments.

### Pathogen apathy from constitutive SKN-1 activation is a defect in serotonin signaling

The GO-term enrichment among the genes that differentially respond to PA14 based on the presence of the *skn-1gf* allele was intriguing and we noted a distinct difference in the genotype-dependent response of genes with roles in neuronal function (**Figure 5A**). Moreover, genes involved in serotonin signaling (**Figure 5B**) were particularly sensitive to both genetic and dietary conditions (**Figure 5C** and Figure S5A). With this in mind, we tested whether treatment with the monoamine neurotransmitter serotonin (5-hydroxytryptamine, 5-HT) could influence pathogen leaving behavior. Strikingly, treatment of *skn-1gf* animals with serotonin from the earliest larval stage (L1) significantly reversed the pathogen leaving behavior without changing the leaving behavior of WT animals, as compared to vehicle treated controls (**Figure 5D**). This result was specific to serotonin (5-HT), as treatment with 5-hydroxytryptophan (5-HTP), the rate-limiting precursor of serotonin biosynthesis (Figure S5B) did not reverse apathy for leaving the pathogen environment observed in mock-treated *skn-1gf* animals. Similarly, treatment with other neurotransmitters, including dopamine (Figure S5C) and octopamine (Figure S5D), was unable to alter pathogen leaving behaviors which suggests that serotonin is specifically limiting in *skn-1gf* mutant animals. Serotonin availability is regulated at multiple levels (**Figure 5B**) including reuptake, which effectively recycles serotonin previously released and as such, we tested what would happen in animals treated with fluoxetine, a selective serotonin reuptake inhibitor (SSRI) and found that *skn-1gf* mutants displayed enhanced sensitivity to fluoxetine, resulting in a significant delay in the development into adulthood as compared to WT animals (**Figure 5E**). Collectively, these data suggest that *skn-1gf* mutants have diminished serotonin availability that influences behavioral responses to PA14.

**Figure 5.**
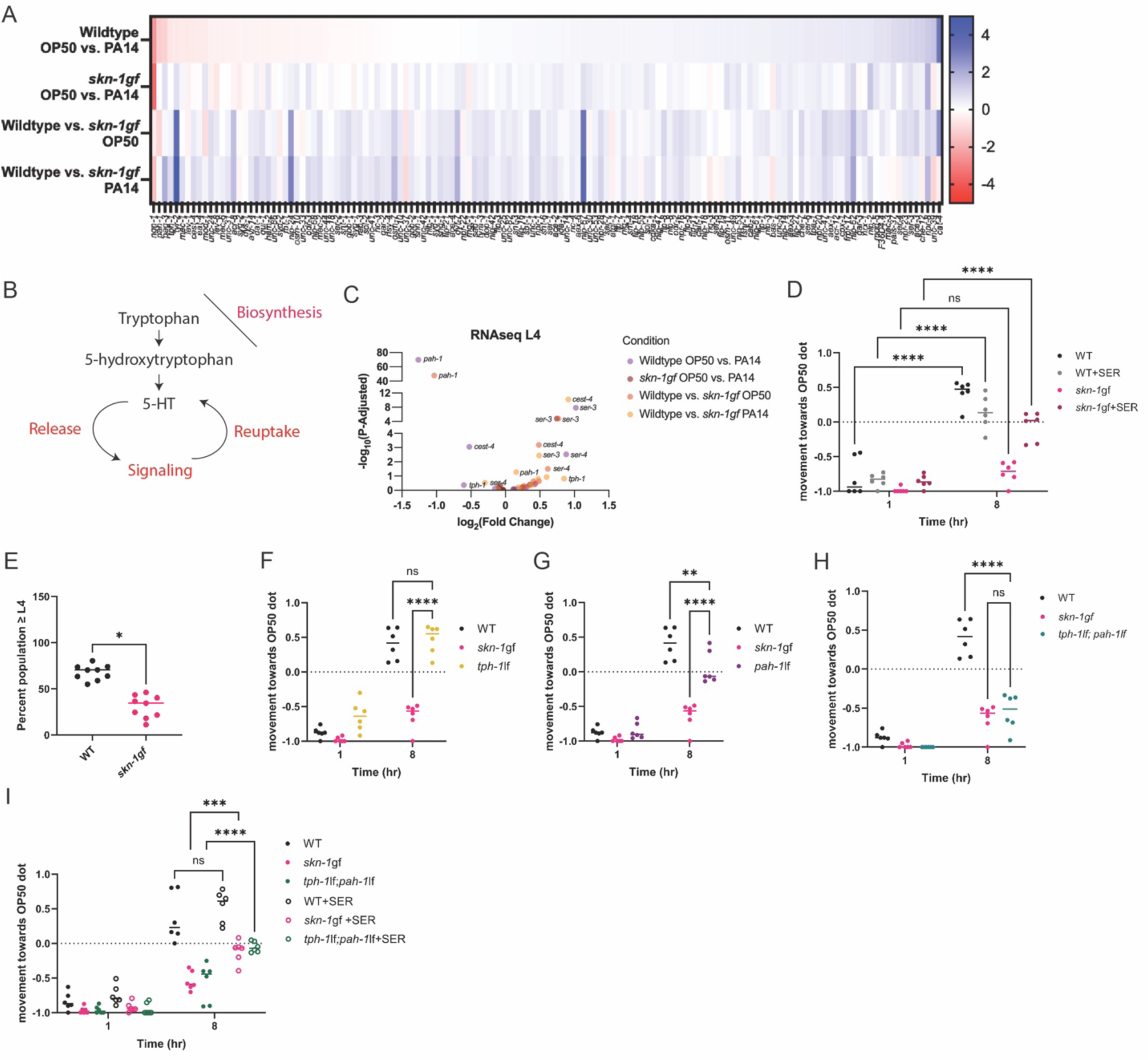
Pathogen apathy is a serotonin signaling defect. (**A**) Heat map of neuronal enrichment genes differentially expressed upon PA14 exposure between WT and *skn-1gf* animals, in comparison to OP50. (**B**) Representation of serotonin signaling pathway including biosynthesis, uptake, and reuptake. (**C**) Selective representation of differentially expressed genes involved in serotonin signaling between WT and *skn-1gf* upon PA14 and OP50 exposure. (**D**) Serotonin supplementation alleviates the apathy response of *skn-1gf* animals. (**E**) *skn-1gf* animals display enhanced sensitivity to treatment with the selective serotonin reuptake inhibitor fluoxetine. Pathogen leaving behavior of (**F**) *tph-1lf* (**G**) *pah-1lf* is similar to WT. (**H**) *tph-1lf;pah-1lf* double mutants display pathogen apathy similar to *skn-1gf* mutants, this apathy is alleviated with supplementation of serotonin (**I**). Each of the food choice assay comprised of N≥3 and analyzed via two-way ANOVA test; **(p<0.01) ***(p<0.001) ****(p<0.0001).

To confirm that the apathy-like behavioral response to PA14 was due to a lack of serotonin and not another outcome of constitutive SKN-1 activation, we next tested mutants in the serotonin biosynthesis pathway for pathogen leaving responses. We tested loss-of-function (lf) mutations in the tryptophan hydroxylase*, tph-1lf* that regulates serotonin biosynthesis in neurons [40]. Surprisingly, *tph-1* mutants displayed an unremarkable leaving response from PA14 as compared to WT (**Figure 5F**). Recently, the phenylalanine hydroxylase, *pah-1lf,* was discovered to regulate non-neuronal biosynthesis of serotonin in non-neuronal tissues [41]. We tested two alleles of *pah-1lf* (**Figure 5G** and Figure S5E), but neither significantly influenced pathogen leaving behaviors. Recalling that *skn-1gf* expression in neurons and intestine alone could not fully recapitulate the pathogen-leaving behaviors of the *skn-1gf* mutant we next tested a *tph-1lf pah-1l*f double mutant animals that strikingly resembled the *skn-1gf* mutant pathogen apathy behavior (**Figure 5H**). Similar to animals with constitutive activation of SKN-1, this failure of *tph-1lf pah-1l*f double mutant animals to leave a pathogenic environment is reversed with 5-HT treatment (**Figure 5I**). Taken together, these data reveal the serotonin-dependent influence of pathogen apathy has contributions from both neuronal and non-neuronal tissues.

### Serotonin depletion limits pathogen resistance

The impaired pathogen leaving behavior observed in the *skn-1gf* mutants is a failure in one of the earliest responses to PA14 exposure that normally occurs within one hour of encountering toxins produced by the bacteria (**Figure 1**). Although serotonin signaling from the nervous system is important for host-pathogen behaviors [15], a role for non-neuronal serotonin [41] in response to SKN-1 activity in ASI neurons is not known. As such, we examined whether conditions that influence pathogen apathy behavior also influence pathogen resistance in the “fast kill” assay, which is a response dependent on the presence of virulence factors that also contribute to pathogen avoidance behaviors. We previously reported that *skn-1gf* mutants display enhance pathogen resistance (Epr) when compared to WT animals [8] and surprisingly treatment with 5-HT resulted in a further enhancement of the Epr phenotype in the *skn-1gf* mutant but had negligible effects on WT animals survival in the fast kill paradigm (**Figure 6A** and Table S4). We also examined the serotonin biosynthesis mutants in the fast kill assay and discovered that the *tph-1lf pah-1lf* double mutant animals display increased sensitivity to PA14 exposure (**Figure 6B**), as compared to either single mutant. These data indicate a role for both neuronal and non-neuronal serotonin signaling in host behavioral responses to pathogens and survival from pathogen infection. As such, pathogen leaving behavior and pathogen resistance are serotonin-sensitive physiological responses that are influenced by SKN-1 activity in both neuronal and non-neuronal tissues.

**Figure 6.**
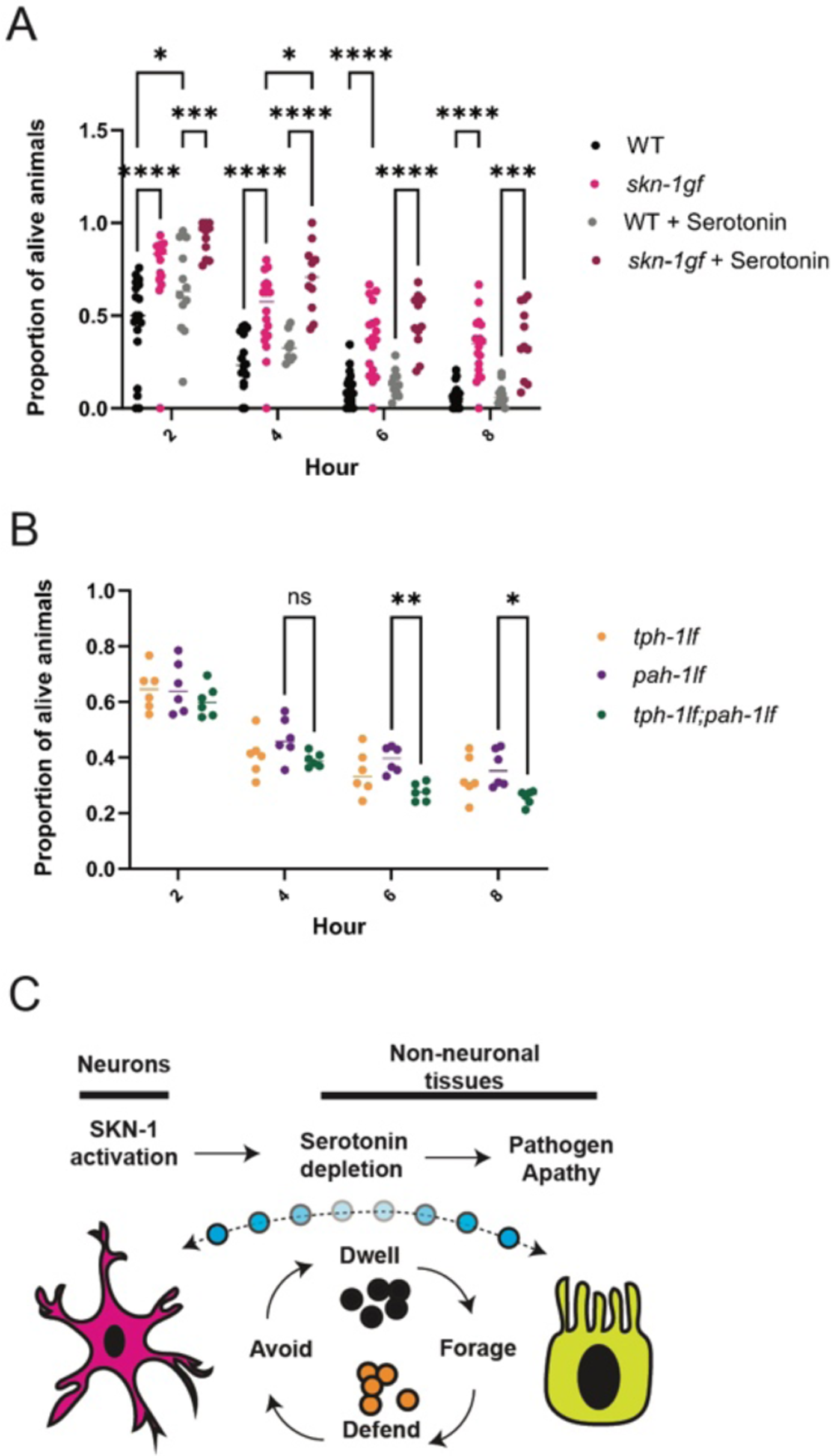
Serotonin depletion limits pathogen resistance. (**A**) Survival analysis of WT and *skn-1gf* animals with and without serotonin supplementation treatment on PA14 fast kill assay plates. (**B**) Survival analysis of *tph-1lf*, *pah-1lf*, and *tph-1lf;pah-1lf* on PA14-seeded fast kill assay plates. (**C**) Cartoon model of the impact of constitutive activation of SKN-1 on serotonin signaling and pathogen apathy. Each of the fast kill assay comprised of N≥3 and analyzed via two-way ANOVA test; **(p<0.01) ***(p<0.001) ****(p<0.0001).

Taken together, these data reveal a consequence of constitutive SKN-1 activation is the depletion of available serotonin that leads to a defect in pathogen leaving behavior. This phenotype is critical for survival in response to exposure to a bacterial pathogen and connects SKN-1 functions and serotonin signaling in cytoprotective defenses to animal decisions on food-dependent behaviors like whether to forage, avoid, or dwell (**Figure 6C**). Collectively, our study leads to a new model to understand how the inability to turnoff of cytoprotective pathways results in pleiotropic consequences that are relevant to health across the lifespan.

## DISCUSSION

Host-pathogen interactions are complex and an animal’s ability to sense and respond to potentially toxic components in the diet is essential for long-term health, and often times, survival. In their natural environment, *C. elegans* are exposed to numerous bacteria, some of which are potentially pathogenic [42, 43]. A proper immune response is required to survive the immediate toxins produced by pathogenic bacteria [44], but the worm must also be able to escape and avoid the pathogen or potential pathogen colonization in the gut, which risks further long-term damage and eventual death. Our work demonstrates how SKN-1 activity is critical for the earliest behaviors post-exposure to a pathogen, and as such, have developed a new model to test how the gut and brain coordinate to ensure an appropriate organism-level response is executed when an animal encounters a potentially toxic environment.

Although not fully understood, several pathogens, like PA14, stimulate attraction behaviors through the olfactory system in *C. elegans,* leading to a preferential association with these microbes over the standard OP50 *E. coli*/B diet [15, 22]. Although *skn-1gf* mutants behave similarly to WT animals in their initial choice of PA14 over OP50, they fail to evoke normal pathogen leaving behaviors, thus appearing apathetic to the pathogenic environment. Two prominent models of leaving behavior link a sophisticated coordination of gut-brain signaling initiated from sensory neurons [17] or intestinal bloating caused by pathogen infection that send cues to the nervous system to drive pathogen avoidance [19]. Our endogenous SKN-1-GFP reporters confirm the steady-state presence of SKN-1 exclusively in ASI neurons [32], which is an essential neuron that is required for proper transgenerational inheritance of pathogen avoidance [45], however, expression of *skn-1gf* only in ASI was not sufficient to influence pathogen leaving behavior, indicating the necessity of other neurons to elicit this response. This idea is supported by specifically degrading activated SKN-1 throughout the nervous system, restores pathogen leaving behaviors. Moreover, our discovery of the precocious nuclear accumulation of SKN-1 within intestinal nuclei, upon exposure to PA14, further connects the brain-gut axis model of SKN-1 activity to pathogen related behaviors and survival.

Our first evidence that constitutive SKN-1 activation resulted in a differential response to pathogens, as compared to animals that maintain the capacity to turn SKN-1 off, was in the distinct transcriptional response to PA14, which included changes in multiple serotonin signaling pathway genes. 5-HT can potently stimulate fatty acid B-oxidation, which results in fat loss in the intestine and animals with constitutive activation of SKN-1 deplete intestinal lipids in an age-dependent manner [8, 46]. Previous work has demonstrated that serotonin signaling suppresses the innate immune response, limiting the rate of pathogen clearance, and that serotonin released from sensory neurons modulates the immune system in response to changes in the animal’s external environment [47]. SKN-1 is a key regulator of environmental stress and constitutive activation, specifically in neurons, is required for several pleiotropic responses to the inability to turn off this cytoprotective response [32]. Perhaps most important is our finding that serotonin supplementation, in the context of constitutively active SKN-1, results in enhanced pathogen resistance indicating that the loss of available serotonin resulting from activated SKN-1 limits organismal capacity to fight off infection.

Our finding that the apathy behaviors observed in the *skn-1gf* mutants can be reversed by treatment with serotonin is intriguing as serotonin signaling is associated with food responses across animals, but also apathy-behaviors in humans [48, 49]. The apathy observed in *skn-1gf* mutants is phenocopied only when serotonin synthesis is disrupted in both neuronal and non-neuronal cell types, suggesting that at least for basal-levels of activity, a certain degree of compensation can occur. In many patients taking selective serotonin reuptake inhibitors (SSRIs), a common treatment for depression, outcomes associated with loss of motivation, energy, and lack of curiosity (collectively referred to as apathy) are commonly experienced [49]. Our discovery reveals that *skn-1gf* mutant animals exhibit a synthetic interaction with serotonin depletion by SSRI treatment that results in developmental arrest and mimicking the worsened outcomes of individuals starting or increasing SSRI treatment, especially if they have apathy - which is an often misdiagnosed condition mistaken for depression. With this in mind, our work suggests that NRF2 activation could be a facile biomarker to measure when considering treatments that manipulate serotonin signaling.

Nuclear factor erythroid 2-related factor 2 (Nrf2), the human homologue of SKN-1, is known for its pivotal role in inducing antioxidant stress and regulating inflammatory responses. It plays a crucial role in the progression of intestinal fibrosis and carcinogenesis associated with inflammatory bowel diseases (IBD) [50]. The inflammatory response exhibited upon the onset of Ulcerative colitis (a form of IBD) leads to the dysregulation of the immune system, and constitutive activation of Nrf2 has been demonstrated to exacerbate cases of acute colitis to carcinoma [50, 51]. Serotonin synthesized mainly in the GI tract is involved in a vast array of biological processes revolving around the gut-brain axis. It also plays a role in regulating inflammation, as increased inflammation in IBD lowered serotonin reuptake [52]. Further, apathy is associated with polymorphisms in serotonin uptake transporter genes that affect their functionality [53, 54] and serotonin reuptake inhibitor (SSRI) treatments offered to patients with depression often display higher cases of apathy [55]. As such, physiological responses revealed in this study of *C. elegans* could inform and model human disease states at the intersection of diet, behaviors, and immunity axes.

## SUPPLEMENTAL FIGURES

**Supplemental Figure S1.**
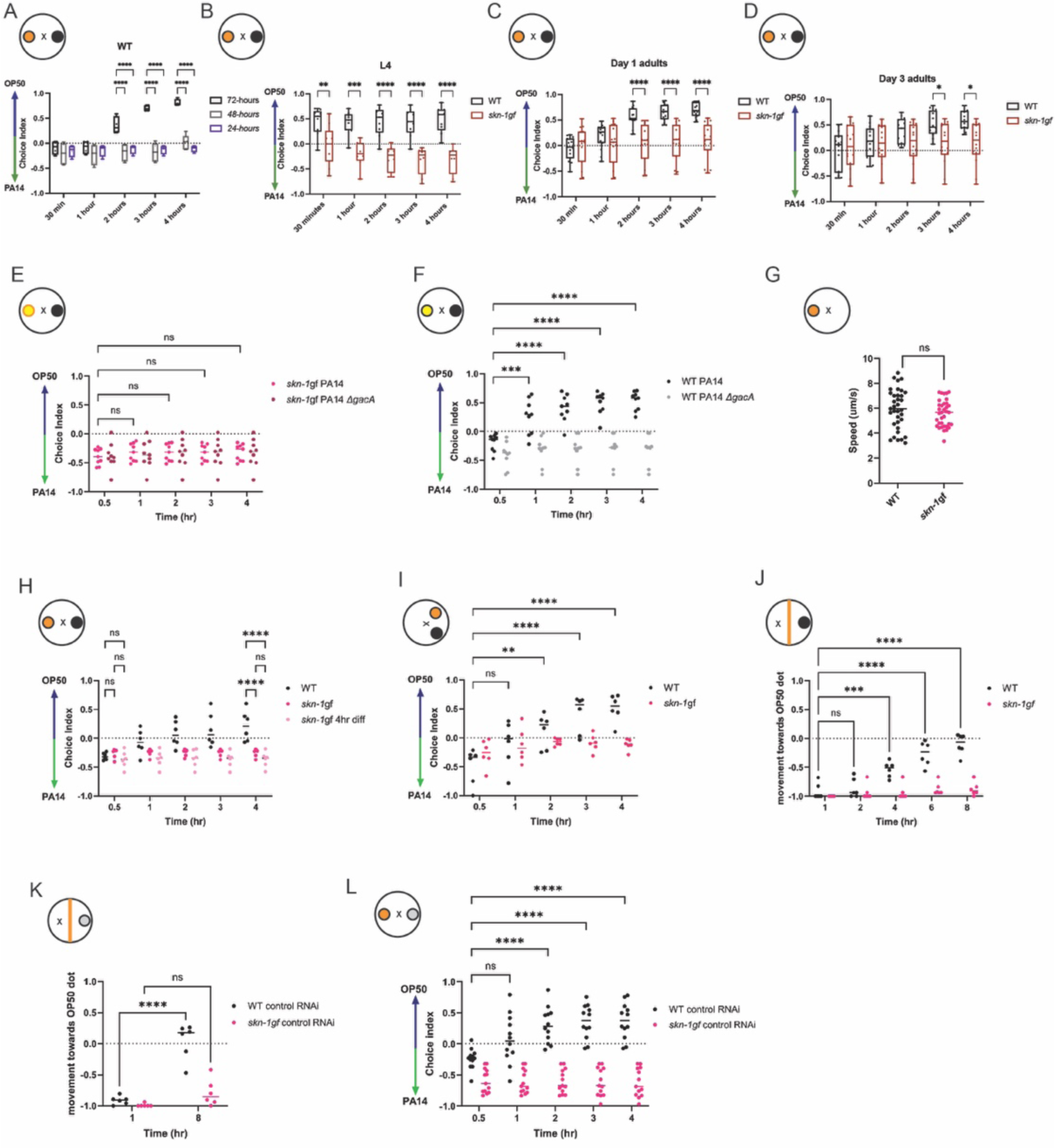
Factors that affect SKN-1gf driven pathogen apathy. (**A**) Pathogen response in WT animals across various PA14 cultures: 72 hr (37°C for 24 hours, followed by 48 hours at 25°C), 48 hr (37°C for 24 hours, followed by 24 hours at 25°C), and 24 hr at 37°C. (**B-D**) Apathy response in WT and *skn-1gf* worms across developmental stages (L4, Day 1 adulthood, and Day 3 adulthood). (**E-F**) Apathy response of WT and *skn-1gf* animals for non-virulent PA14 *ΔgacA* mutants. (**G**) Similar movement speeds of WT and *skn-1gf* animals towards the first chosen diet. (**H**) 4hr difference in developmental timing between WT and *skn-1gf* animal does not alter pathogen apathy response in *skn-1gf* animals. (**I**) Changes in the distance between the two diets does not alter the pathogen apathy response in WT and *skn-1gf* animals. (**J**) “Forced” choice assay, where *skn-1gf* mutant animals reach the PA14 line within 60 min, similar to WT animals but remain on PA14, whereas the WT animals move toward the *E. coli* OP50 option over the 8-hour time course. (**K-L**) *skn-1gf* display apathy in both choice assays for PA14 when first grown on control RNAi (*E. coli* HT115) until L4 stage. Each of the food choice assay comprised of N≥3 and analyzed via two-way ANOVA test; **(p<0.01) ***(p<0.001) ****(p<0.0001) and the movement analyses via unpaired t test; **(p<0.01) ***(p<0.001) ****(p<0.0001)

**Supplemental Figure S2.**
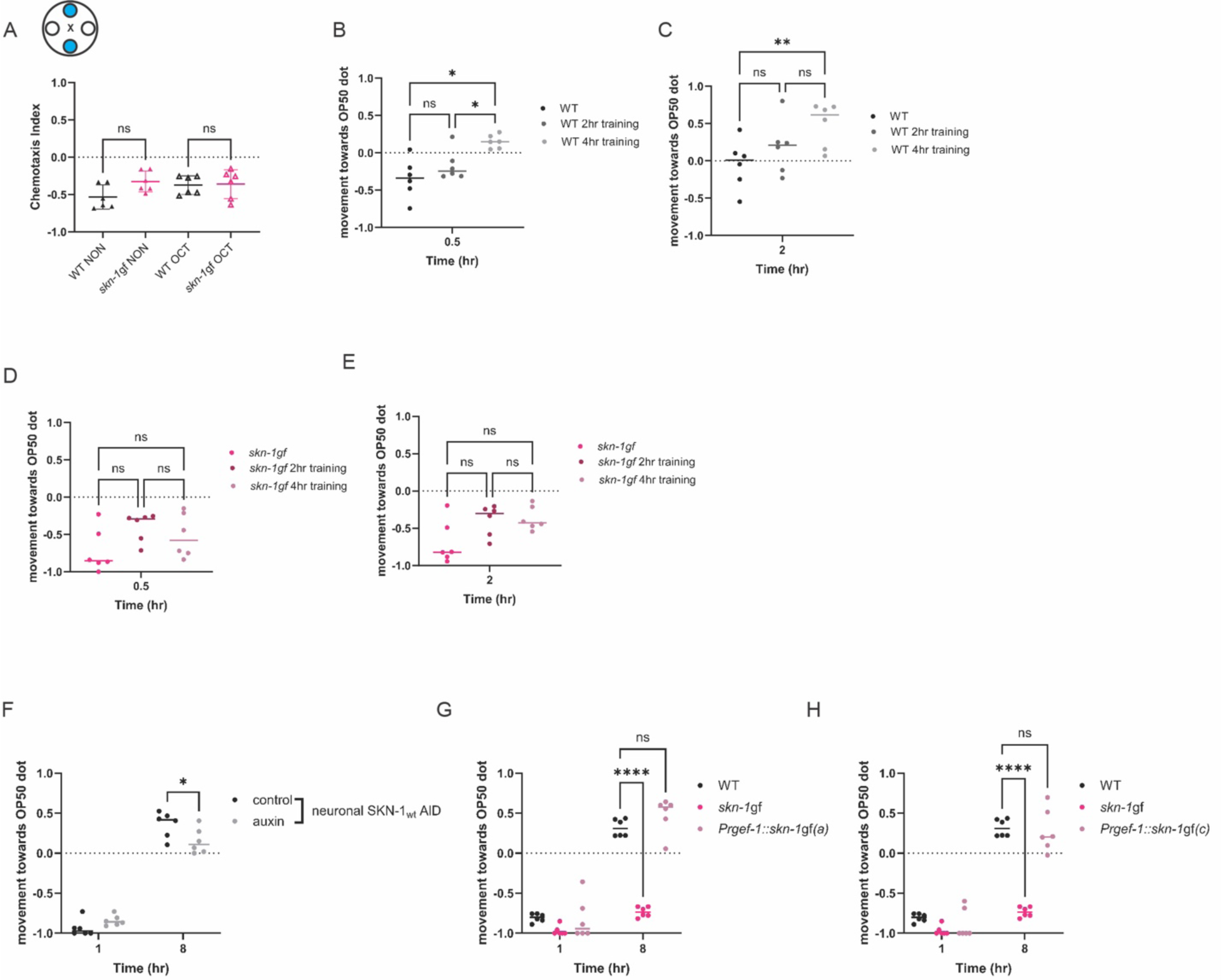
Training and neuronal responses of *skn-1gf*. (**A**) Chemotaxis away from the chemo repellants [octanol (OCT), and nonanone (NON)] is similar between WT and skn-1gf mutants. Pathogen leaving responses in WT (**B-C**) and *skn-1gf* (**D-E**) worms post-pathogen training (0hr, 2hr, and 4hr exposure). (**F**) Pan-neuronal (*rgef-1p*) degradation of SKN-1wt does not alter pathogen leaving behavior. Pan-neuronal (*rgef-1p*) expression of SKN-1gf isoform a (**G**) or isoform c (**H**) in the neurons does not elicit an apathy response similar to *skn-1gf* worms. Each of the food choice assay comprised of N≥3 and analyzed via two-way ANOVA test; **(p<0.01) ***(p<0.001) ****(p<0.0001).

**Supplemental Figure S3.**
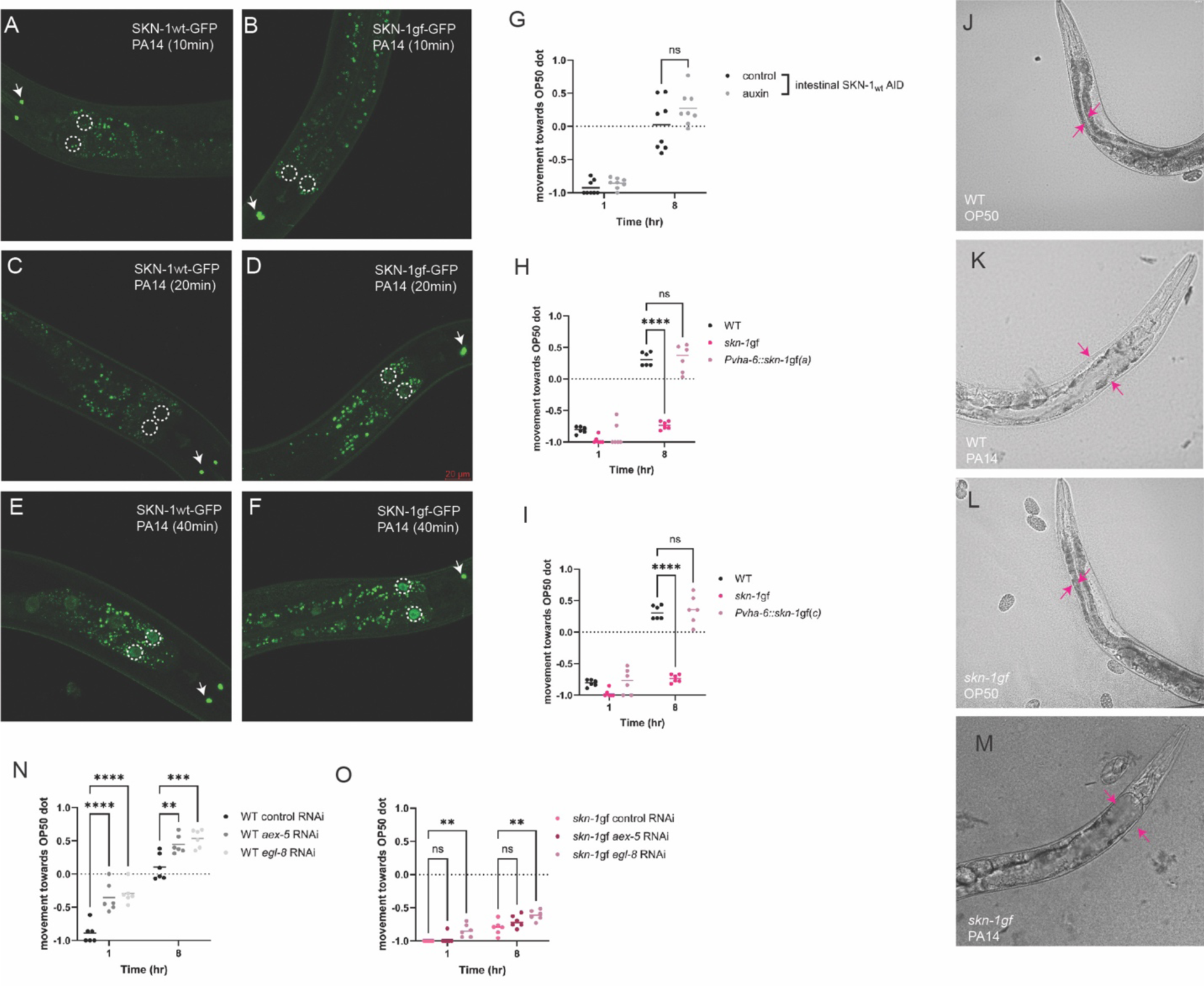
Nuclear accumulation of SKN-1 and intestinal bloating in response to PA14. (**A-F**) Intestinal accumulation and stabilization of SKN-1gf-GFP and SKN-1wt-GFP upon PA14 exposure in comparison to OP50 (time points: 10 mins, 20 mins, and 40 mins); the first two intestinal nuclei of each worm are outlined by white-dotted circles. (**G**) Intestinal-specific (*ges-1p*) degradation of SKN-1wt does not alter pathogen leaving behavior in SKN-1wt mutants. Intestinal-specific (*ges-1p*) expression of SKN-1gf isoforms a (**H**) or c (**I**) is not sufficient to elicit the pathogen apathy behavior as observed in *skn-1gf* worms. (**J-M**) DIC images of intestinal lumen in (**J,K**) WT and (**L,M**) *skn-1gf* worms post-pathogen exposure (24hrs of exposure to PA14). (**N-O**) RNAi of *aex-5* or *egl-8* in WT animals causes enhanced pathogen leaving even in the absence of PA14, however only *egl-8* RNAi was capable of marginally inducing leaving behavior of the *skn-1gf* mutant. Each of the nuclear accumulation time points comprised of N≥3 and analyzed via two-way ANOVA test; **(p<0.01) ***(p<0.001) ****(p<0.0001). All experiments were conducted at 25°C.

**Supplemental Figure S4.**
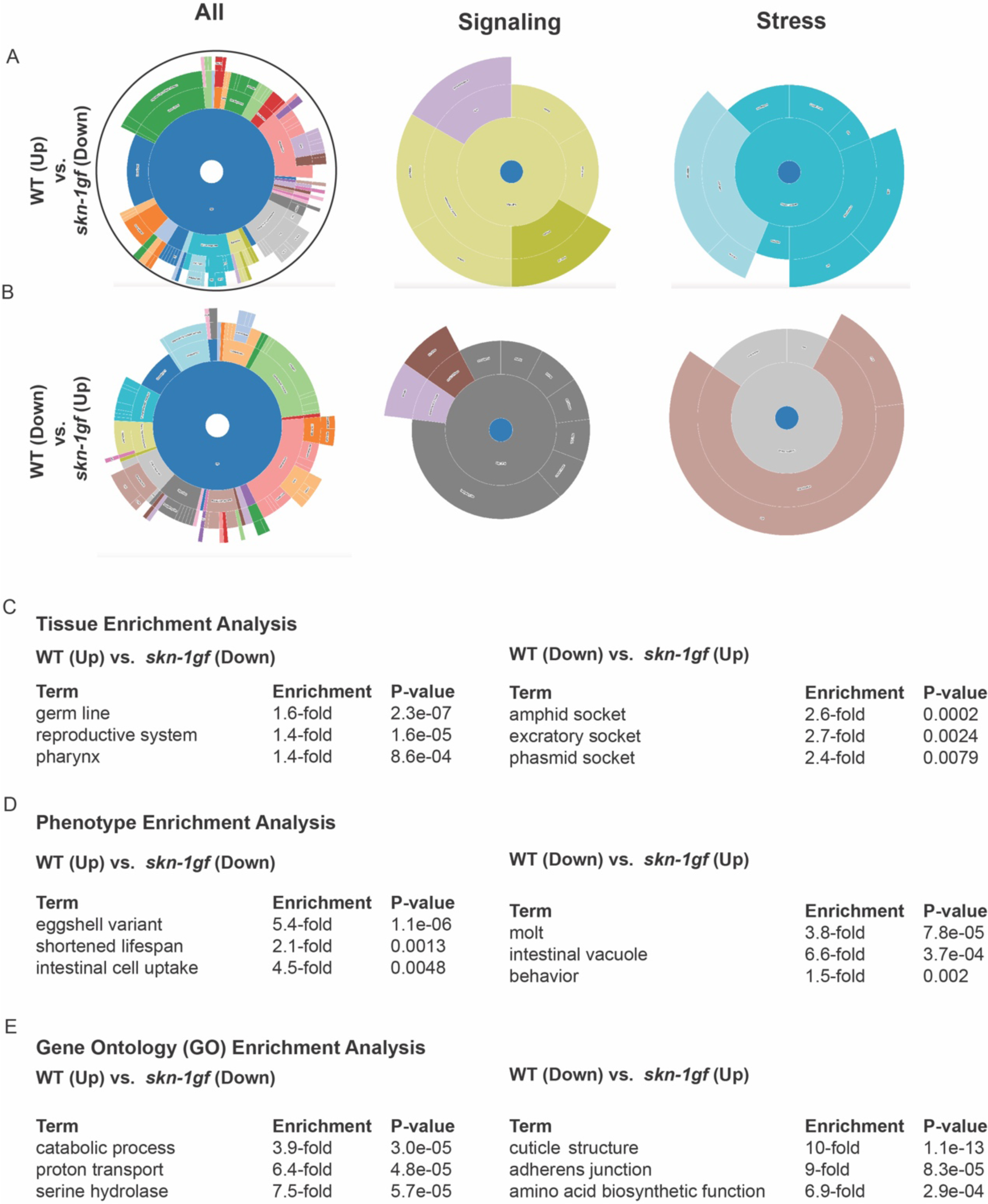
Gene set enrichment analysis (GSEA) of context specific transcriptional signature of PA14 exposure. Cartoon visualization of differences in gene set enrichment analysis from RNAseq data analyzed in WormCat [56] of PA14 responsive genes that are (**A**) up-regulated in WT but down-regulated in *skn-1gf* or (**B**) down-regulated in WT but up-regulated *in skn-1gf* (see Supplemental Table S3). Gene set enrichment analysis (GSEA) tools in WormBase were used for the RNAseq data in Supplemental Table S3 to identify (**C**) tissue enrichment, (**D**) phenotype enrichment, and (**E**) gene ontology (GO) term enrichment of transcripts that differentially respond to PA14 in WT and *skn-1gf* animals.

**Supplemental Figure S5.**
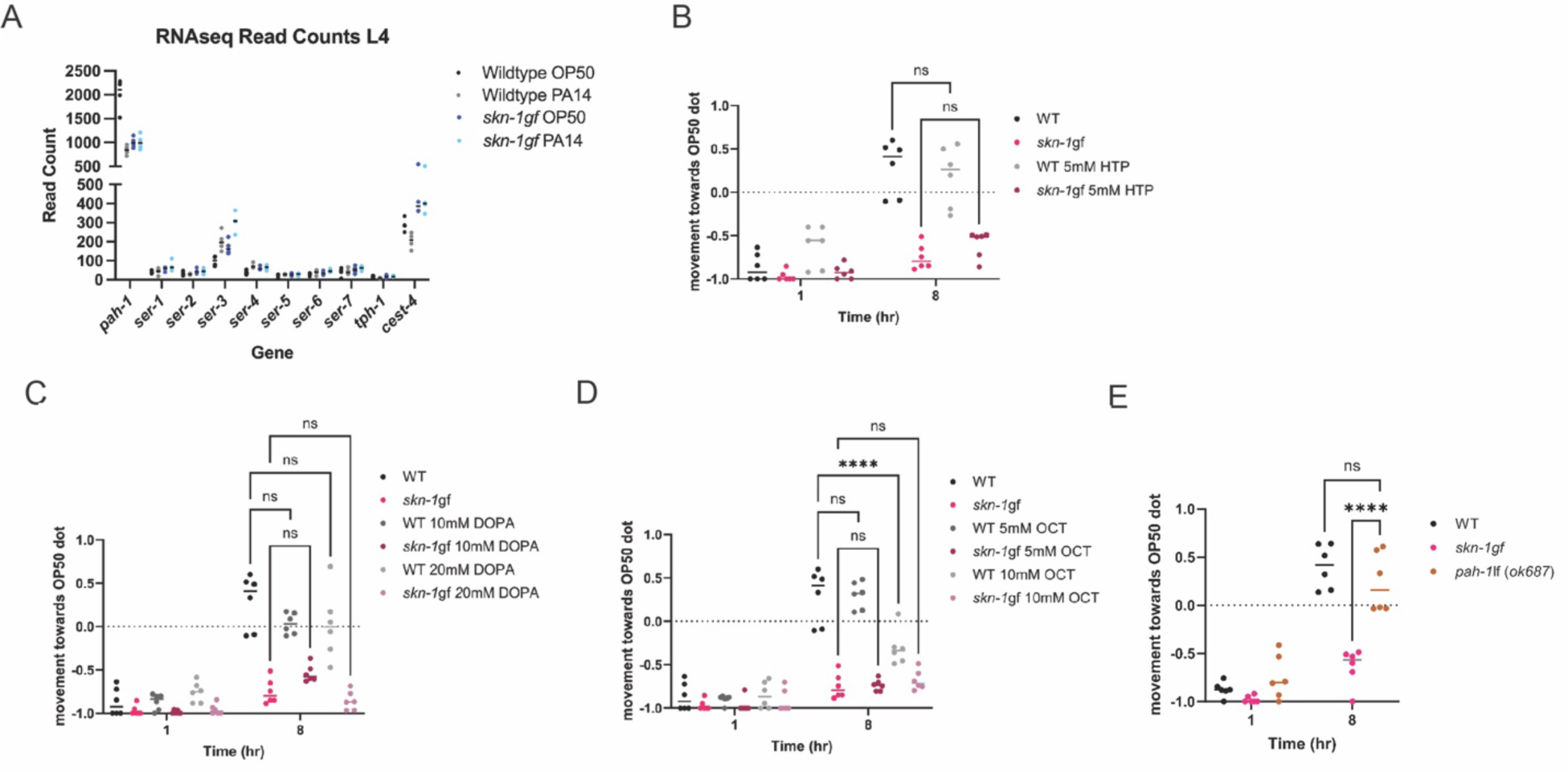
Pathogen apathy with constitutive SKN-1 is associated exclusively with serotonin. (**A**) The GO-terms enrichment analysis of serotonin pathway genes that differentially respond to genetic and dietary conditions in WT and *skn-1gf* animal upon exposure to PA14, in comparison to OP50. Supplementation of (**B**) 5mM 5-HTP, (**C**) 10-20mM dopamine, (**D**) or 5-10mM octopamine does not alleviate the pathogen apathy response in *skn-1gf* animals. (**F**) No apathy response observed in the serotonin pathway mutant *pah-1l*f(*ok687*). Each of the food choice assay comprised of N≥3 and analyzed via two-way ANOVA test; **(p<0.01) ***(p<0.001) ****(p<0.0001). All experiments were conducted at 25°C

## SUPPLEMENTAL TABLES

Table S1. Behavioral responses to PA14

Table S2. SKN-1 nuclear activation response

Table S3. Genotype-specific transcriptional responses to PA14 exposure at the L4 stage.

Table S4. Pathogen survival data

## MATERIALS AND METHODS

### *C. elegans* strains and maintenance

All worms were grown at 20°C on 6 cm Nematode Growth Medium (NGM) agar plates with streptomycin and seeded with 250μL *E.coli* strain OP50-1, unless otherwise noted. The following strains were used: WT – N2 Bristol (reference strain), SPC227 - *skn-1gf(lax188),* SPC2002 - *skn-1wt-GFP,* SPC2003 - *skn-1gf-GFP,* SPC602 *- skn-1wt-AID; rgef-1p::TIR1,* SPC603 - *skn-1wt-AID; ges-1p::TIR1,* SPC2048 - *skn-1gf-AID; rgef-1p::TIR1,* SPC2047 - *skn-1gf-AID; ges-1p::TIR1,* SPC2065 - *rgef-1p::skn-1gf-isoformA,* SPC2064 - *rgef-1p::skn-1gf-isoformC,* SPC2058 - *vha- 6p::skn-1gf-isoformA,* SPC2057 - *vha-6p::skn-1gf-isoformC,* SPC2108 - *vha-6p::skn-1gf-isoformA;vha-6p::skn-1gf-isoformC,* SPC2068 - *gpa-4p::skn-1gf-isoformA,* SPC2067 - *gpa-4p::skn-1gf-isoformC,* MT15434 - *tph-1(mg280*), LC74 - *pah-1(ok687)*, PHX3596 - *tph-1(mg280) pah-1(syb3596),* PHX3601 - *pah-1(syb3601)*

For all experiments, eggs were collected from day 1 adult animals by hypochlorite treatment and allowed to hatch overnight for L1 synchronization. For testing on day 3 adults, worms had to be washed to new plates every day beginning at day 1 of adulthood, in order to separate worms from progeny. M9 + 0.01% Triton X-100 (M9T) was used to wash the worms from the plate into microcentrifuge tubes. Worms were allowed to gravity settle and the supernatant containing the progeny was removed without disturbing the adult pellet. This was repeated three times, and worms were dropped onto fresh NGM plates.

### Bacterial strain and maintenance

*E. coli,* strains OP50-1 and HT115 and *P. aeruginosa,* strain PA14 and PA14Δ*gacA* were used. All strains were streaked onto LB agar plates, with different antibiotic conditions for each strain. OP50 was streaked on plates with streptomycin, HT115 on plates with ampicillin, PA14 and PA14Δ*gacA* on plates without antibiotics. OP50 that was used for preference plates were streaked on LB plates without antibiotics to match PA14. Following streaking, plates were grown overnight at 37°C and either used the next day or stored at 4°C for no longer than a week.

#### RNAi experiments

RNAi treatment was performed as previously mentioned [57]. Briefly, *E. coli* HT115 bacteria carrying specific double stranded RNA-expression plasmids were seeded on NGM plates containing 5mM isopropyl-b-D-thiogalactoside (IPTG) and 50mgml-1 carbenicillin. RNAi were induced at room temperature for 24 h. Synchronized L1 animals were added onto RNAi plates to knockdown indicated genes. L4 animals were subjected to food choice assays, and choice index was measured.

#### Fast Kill Assays

Fast Kill Assay were performed in 3.5 cm peptone glucose media plates (1% Bacto-Peptone; 1% NaCl; 1% Glucose; 1.7% Bacto-Agar) containing 0.15M sorbitol. PA14 cultures were grown overnight at 37°C for 14-15 hours. 5ul of the overnight culture was spread over the PGM +sorbitol plates using a spreader made from an open loop tipped glass pasture pipette. The plates were incubated at 37°C for 24 hours and then at 25°C overnight. 30-40 L4 animals were placed on each plate. Assay was performed at 25°C. Survival of animals was plotted over a period of 8 hrs with intervals of 2 hrs (0 hr, 2hr, 4hr, 6hr, 8hr) [58, 59].

### Behavioral analyses

#### Two-choice preference assay

Preference plates were made by using the same NGM plates for growth, except 0.25% peptone is replaced by 0.35% peptone, and no antibiotics are used. The night before seeding, a single colony from each bacteria that is to be used in the preference assay was picked into 3mL of LB and grown in a 37°C shaker overnight. The following day, the OD600 was measured by diluting 100μL of bacteria with 900μL of LB and multiplying the OD600 readout by ten. The bacteria were then diluted to an OD600 of 1.0. Two spots were marked equidistant from the center, yet not too close to the edge, and 15μL of diluted bacteria was placed on the marked spots. Plates were allowed to dry and then transferred to the 37°C for 24 hours, followed by the 25°C incubator for 48 hours, unless otherwise noted.

On the day of the assay, the plates were removed from the 25°C incubator to reach room temperature. Worms were collected from NGM growth plates in M9 buffer and further subjected to three washes in M9T buffer. Finally, the worms were placed onto the assay plate and recorded.

#### Forced choice assay [60]

Plates were made of similar composition as the two choice preference assay plates. A single colony from each *Pseudomonas aeruginosa* and *E. coli* bacteria were inoculated into 3mL of LB for overnight primary culture at 37°C. The OD600 of each of the primary cultures was measured following which the cultures were diluted to a OD600 of 1.0. A drop of 15μL bacterial culture was seeded onto NGM plates (without streptomycin) equidistant from the center to the point where the animals (worms) were being dropped (not too close to the edge). The complimentary bacterial culture was stamped on the plates diametrically using an “L” tipped glass pasture pipette. The plates were dried and transferred to the 37°C for 24 hours, followed by the 25°C incubation for 48 hours, unless noted otherwise.

On the day of the assay, the plates were removed from the 25°C incubator to reach room temperature. Worms were collected from NGM growth plates in M9 buffer and further subjected to three washes in M9T buffer. Finally, the worms were placed onto the assay plate and recorded.

#### Chemoattraction and chemorepulsion

Chemotaxis assays were performed as previously described [61]. Briefly, the underside of a 5 cm unseeded NGM plates were divided into 4 equal quadrants. A circle of 0.5 cm radius was marked around the center. Diagonally opposite quadrants were marked either for the test odorant or as the vector control, ethanol. The sites where the odorants or Ethanol were to be spotted were equidistant from the center and each other. Equal volumes of the test odorant and 0.5 M azide or control and 0.5 M azide were mixed to make working solutions. Young adult worms were washed and resuspended in 100 ul of M9 buffer. 50-100 worms were added onto the center of the plate. Immediately after which 2 μl of the test solution and control solution were added to the respective sites in each quadrant. Once the odorant drops were absorbed in the agar the lids were placed and plates were inverted. After 60 min, the assay plates were placed in a 4°C. Only the assay plates required to be assessed and counted were removed from 4°C. The number of worms in each quadrant that crossed the center were recorded. For each odorant set replicates were in an order of (n=3&N=3). The chemotaxis index was measured as follows: Chemotaxis Index = (# Worms in Both Test Quadrants - # Worms in Both Control Quadrants) / (Total # of Scored Worms). +1.0 score indicated maximal attraction towards the compound and −1 indicated maximal repulsion.

### Pharmacological treatments

Serotonin hydrochloride; 5-HT (Sigma, H9523) powder was dissolved in MilliQ to a working stock concentration of 0.1M and added to NGM plates seeded with *E. coli* OP50 to a final plate concentration of 5mM [62]. Plates were allowed to equilibrate, and synchronized L1-stage animals were added onto 5-HT-treated plates. At the L4 stage, worms were then subjected to the indicated food choice assays and physiological assays like fast kill and subjected to the visualization of pathogen mediated localization of SKN-1.

Dopamine Hydrochloride; DOPA (Sigma, H8502) powder was dissolved in MilliQ to a working stock concentration of 1M and added to NGM plates seeded with *E. coli* OP50 to a final plate concentration of 10mM and 20mM [63]. Plates were allowed to equilibrate, and synchronized L1-stage animals were added onto dopamine-treated plates. At the L4 stage, worms were then subjected to the indicated food choice assays.

5-hydroxytryptophan; 5-HTP (Sigma, H9772) powder was dissolved into a working stock concentration of 0.1M and added to NGM plates seeded with *E. coli* OP50 to a final plate concentration of 5mM [64]. Plates were allowed to equilibrate, and synchronized L1-stage animals were added onto 5-HTP-treated plates. At the L4 stage, worms were then subjected to the indicated food choice assays.

Octopamine Hydrochloride; OCT (Sigma, O0250) powder was dissolved in MilliQ and added to NGM plates to make the final concentrations of 10mM and 5mM prior to plates were seeded with *E. coli* OP50 [65]. Synchronized L1-stage animals were added onto Octopamine treated plates. At the L4 stage, worms were then subjected to the indicated food choice assays.

Fluoxetine Hydrochloride (Sigma, PHR1394) powder was dissolved in MilliQ and added to NGM plates seeded with *E. coli* OP50 to make the final concentrations of 145 µM (0.5mg/ml) [66]. Synchronized L4-stage animals were added onto Fluoxetine treated plates to assess survival percentages.

### Crawling Assays

Worms were egg prepped and eggs were allowed to hatch overnight for a synchronous L1 population. The next day, worms were dropped onto plates seeded with OP50. Worms were then allowed to grow until each time point (48 h post-drop for L4s). Once worms were at the required stage of development, 30-50 worms were washed off of a plate in 50 uL of M9 with a cut and M9+triton coated P1000 tip and dropped onto PA14 plates (as described above).

Pseudomonas (PA14) was prepared with an overnight culture of LB incubating at 37°C. A 15µL drop of PA14 was placed on one end of a standard NGM plate with no streptomycin. Plates were incubated in 37°C for 24 hours, followed by 25°C for 48 hours. L4 worms were placed directly opposite from the bacteria culture. The M9 was allowed to dissipate, time to reach the PA14 food recorded. Crawling was imaged with the MBF Bioscience WormLab microscope and analysis was performed with WormLab version 2022. Worm crawling on the plate was imaged for 1 minute for each condition at 7.5 ms. Values were analyzed under the “Speed” tab. Mean speed values of each individual animal captured overall movement speed for the time interval. Worm crawling was analyzed with the software and only worms that moved for at least 90% of the time were included in the analysis. Irregular speed values were discarded from the dataset using normalization to the overall average of the speed values. Statistical analysis of crawling speed was done via one way ANOVA with multiple t-test.

### RNAseq and analyses

RNAseq analysis was conducted as outlined [43, 67]. Worms were egg prepped and eggs were allowed to hatch overnight for a synchronous L1 population. The next day, L1s were dropped onto seeded NGM plates and allowed to grow 48 h (L4 stage) then exposed to PA14 before collection. Animals were washed 3 times with M9 buffer and frozen in TRI reagent at −80°C until use. Animals were homogenized and RNA extraction was performed via the Zymo Direct-zol RNA Miniprep kit (Cat. #R2052). Qubit™ RNA BR Assay Kit was used to determine RNA concentration. The RNA samples were sequenced and read counts were reported by Novogene. Read counts were then used for differential expression (DE) analysis using the R package DESeq2 created using R version 3.5.2. Statistically significant genes were chosen based on the adjusted p-values that were calculated with the DESeq2 package (p<0.01). Gene Ontology was analyzed using the most recent version of WormCat 2.0 [56].

### Auxin-inducible degradation

For tissue-specific degradation experiments, worms were egg prepped and allowed to hatch overnight for a synchronous L1 population. The next day, worms were dropped onto plates with vehicle 4mM ethanol or 4mM auxin (Sigma Aldrich, I3750) and raised to L4 stage and then moved to experiment plates.

### Imaging

#### Fluorescence

For confocal microscopy (SKN-1-GFP, 491nm), imaging was performed on a Stellaris 5 confocal microscope equipped with a white light laser source and spectral filters, HyD detectors, 63x/1.4 Plan ApoChromat Oil objective, and run on LAS X 4.4.0.24861 software.

#### Intestinal distension

The NGM plates (without streptomycin) were seeded with an overnight culture of *Pseudomonas Aeruginosa* PA14 and allowed to dry. Further the plates were incubated at 37°C for 24 hours, followed by the 25°C incubation for 48 hours. Perfect L4 staged worms (45-48 hrs post L1 starvation) placed onto the PA14 seeded plates and observed over a period of 24 hrs. The images intestinal distension was taken with LAS X software and Leica Thunder Imager flexacam C3 color camera (Magnification-63X)

### Statistics

All statistical analysis were performed using GraphPad Prism version 10.0. Information on specific statistical tests used for particular assay is present within respective figure legends.

## ACKNOWLEDGMENTS

We thank S. Ledgerwood for technical assistance and C.M. Ramos for critical reading of the manuscript. This work was funded by the NIH R01AG058610 and Hevolution Foundation award HF AGE-004 to SPC, T32AG000037 to BAW, and T32AG052374 and F31AG077873 to NLS. We also thank the USC School of Gerontology Imaging Core that is funded in part by the Nathan Shock Center of Excellence P30AG068345. Some strains were provided by the CGC, which is funded by the NIH Office of Research Infrastructure Programs (P40 OD010440). We thank WormBase for database curation and data access.

## Author contributions

Conceptualization: SPC; Methodology: TN, BAW, NLS, JDN, and SPC; Investigation: TN, BAW, NLS, JDN, and SPC; Visualization: TN, BAW, NLS, JDN, and SPC; Supervision: SPC; Writing (original draft): SPC; Writing (reviewing & editing): TN, BAW, NLS, and SPC.

## Data and materials availability

All data are available in the main text or the supplementary materials. RNAseq data is available at the NIH Gene Expression Omnibus (GEO).

